# Male harm suppresses female fitness to affect the dynamics of adaptation and evolutionary rescue

**DOI:** 10.1101/2022.08.19.504524

**Authors:** Miguel Gómez-Llano, Gonçalo S. Faria, Roberto García-Roa, Daniel W.A. Noble, Pau Carazo

**Affiliations:** Department of Biological Sciences, University of Arkansas; School of Biological Sciences, University of East Anglia; Ethology lab, Cavanilles Institute of Biodiversity and Evolutionary Biology, University of Valencia, Valencia, Spain; Division of Ecology and Evolution, Research School of Biology, The Australian National University, Canberra, Australia

**Author notes:** corresponding authors: Miguel Gómez Llano, Department of Biological Sciences, University of Arkansas, Fayetteville, Arkansas, United States; Pau Carazo, Cavanilles Institute of Biodiversity and Evolutionary Biology, University of Valencia, Valencia, Spain.

**Keywords:** sexual conflict, sexual selection, local adaptation, evolutionary rescue

## Abstract

One of the most pressing questions we face as biologists is to understand how climate change will affect the evolutionary dynamics of natural populations and how these dynamics will in turn affect population recovery. Increasing evidence shows that sexual selection favours population viability and local adaptation. However, sexual selection can also foster sexual conflict and drive the evolution of male harm to females. Male harm is extraordinarily widespread and has the potential to suppress female fitness and compromise population growth, yet we currently ignore its net effects across taxa, or its effects on local adaptation and evolutionary rescue. We conducted a comparative meta-analysis to quantify the impact of male harm on female fitness and found an overall negative effect of male harm on female fitness. Negative effects seem to depend on proxies of sexual selection, increasing in species with larger sexual size dimorphism and strong sperm competition. We then developed theoretical models to explore how male harm affects adaptation and evolutionary rescue. We show that, when sexual conflict depends on local adaptation, population decline is reduced, but at the cost of slowing down genetic adaptation. This trade-off suggests that eco-evolutionary feedbacks on sexual conflict can act like a double-edge sword, reducing extinction risk by buffering the demographic costs of climate change, but delaying genetic adaptation. However, variation in the mating system and male harm type can mitigate this trade-off. Our work shows that male harm has widespread negative effects on female fitness and productivity, identifies potential mechanistic factors underlying variability in such costs across taxa, and underscores the importance of male harm on the demographic and evolutionary processes that impact how species adapt to environmental change.

**Impact summary:** For species to persist in the face of climate change, adaptation needs to be fast enough to prevent extinction. If population decline is too abrupt, adaptation will be less likely to promote recovery, leading to extinction. Therefore, numerous studies have sought to determine how species can adapt and escape extinction. Sexual selection can promote genetic adaptation, but often has a by-product, sexual conflict, that promotes adaptations beneficial for one sex and detrimental to the other. Such is the case of male adaptations that increase male reproduction by harming females (male harm). Male harm is widespread and has been shown to decrease female and population productivity in some species, facilitating extinction. Furthermore, there is increasing evidence that the degree of male harm to females depends on environmental changes and how well males are adapted to them. However, we ignore how strong the effects of sexual conflict across taxa are, or how ecological feedback on sexual conflict may affect the rate of adaptation and population recovery. Here, we first conducted a meta-analysis to quantify the effect of male harm on female fitness and show, across taxa, that there is an overall negative effect that seems to be dependent on proxies of sexual selection. Then, we used a series of theoretical models to show that, although eco-evolutionary feedback on sexual conflict can limit population decline, this comes at the cost of slowing down the rate of adaptation and population recovery. Our study suggests that understanding how quick environmental changes affect sexual conflict can increase our understanding of how populations adapt and recover in the face of climate change.

## Introduction

Sexual selection can play a major role in adaptation and evolutionary rescue by promoting genetic adaptation through genic capture and purging the genome of deleterious mutations (Rowe & Houle, 1996; Lorch *et al*., 2003; Parrett & Knell, 2018; Gómez-Llano *et al*., 2020, 2021; Grieshop *et al*., 2021). Furthermore, given that sexual selection is usually stronger on males than females (Janicke & Morrow, 2018; Singh & Punzalan, 2018; Winkler *et al*., 2021), this can be accomplished while minimizing associated demographic costs (Agrawal, 2001; Siller, 2001; Whitlock & Agrawal, 2009; Martinossi-Allibert *et al*., 2018; Grieshop *et al*., 2021). However, sexual selection also tends to favour the evolution of sexual roles (Janicke *et al*., 2016), which foster sexual conflict between the sexes (Arnqvist & Rowe, 2005). In particular, interlocus sexual conflict frequently leads to antagonistic co-evolution, favouring male adaptations that increase male reproductive fitness at the cost of female fitness (Parker, 1979; Arnqvist & Rowe, 2005). Given that population growth is determined to a large extent by female fitness, sexual conflict can reduce population growth and even increase extinction risk (Kokko & Brooks, 2003; Le Galliard *et al*., 2005; Kokko & Rankin, 2006; Rankin *et al*., 2007; Martins *et al*., 2018). Although male harm seems to be extraordinarily diverse and widespread, we currently ignore whether it has net negative consequences for females across taxa, or whether such effects depend on the type of harm. Furthermore, if male harm has widespread demographic effects on populations, these could also impact the process of adaptation and population resilience to environmental change.

To persist in the face of climate change, adaptation must be fast enough to avoid extinction, a process called evolutionary rescue (Gomulkiewicz & Holt, 1995). Evolutionary rescue has an ecological and an evolutionary component. The ecological component is population demography, as population size first declines due to maladaptation, followed by stabilisation and recovery (Gomulkiewicz & Holt, 1995; Carlson *et al*., 2014). Initial population size and decline is a key determinant of evolutionary rescue via bottleneck effects and increased inbreeding, which can further decrease reproduction and/or survival –pushing populations into an extinction vortex (Keller & Waller, 2002; Fox & Reed, 2011; Plesnar-Bielak *et al*., 2012). The evolutionary component is genetic adaptation, as population recovery is achieved by an increase in the frequency of adapted genotypes (Carlson *et al*., 2014). Thus, if a population suffers an abrupt decline or remains at a small population size, the likelihood of extinction increases (Gomulkiewicz & Holt, 1995; Orr & Unckless, 2008; Gomulkiewicz & Houle, 2009; Gomulkiewicz & Shaw, 2013), whereas genetic adaptation marks when recovery can begin and dictates how long population recovery can take. These evolutionary and ecological components set the scene for sexual conflict to play a role in evolutionary rescue, particularly so because recent evidence suggests that male harm, and its impact on populations, depend on the environment. First, male adaptations resulting from sexual conflict are typically condition-dependent, and therefore likely to depend on environmental conditions (Marden & Rollins, 1994; Plaistow & Siva-Jothy, 1996; Fricke *et al*., 2009; Chung *et al*., 2021; Rowe & Rundle, 2021). Second, independent of condition, environmental fluctuations can affect the expression and maintenance of traits involved in male harm, and thus their impact on population viability (Perry & Rowe, 2018; García-Roa *et al*., 2020; Plesnar-Bielak & Lukasiewicz, 2021). This means that males in maladaptive environments will be less capable of harming females, raising intriguing questions about the interplay between environmental change, sexual selection, and evolutionary rescue.

In this study, we aimed to explore whether male harm negatively affects female fitness across taxa, and how ecological feedbacks on the costs of male harm may affect evolutionary rescue in terms of both its ecological (i.e., demographic) and evolutionary (i.e., genetic adaptation) components. To achieve this aim, we first conducted a comparative meta-analysis on studies manipulating male harm levels and measuring the fitness consequences for females. We analysed whether variation in costs of male harm are dependent on different proxies of pre- and post-copulatory sexual selection (i.e., sexual size dimorphism and sperm competition intensity) and the type of harm (i.e., direct and indirect harm). Then, we developed a population genetic model and used numerical simulations to study the effects of male harm on the rate of genetic adaptation and population recovery. We further explored how variation in the mating system modulate the effects of male harm in evolutionary rescue.

## Methods

### 1 Meta-analysis

We conducted a systematic review following the PRISMA protocol (Liberati *et al*., 2009) to look for studies that experimentally manipulated the level of male harm to females and measured its outcome in terms of female fitness. We conducted three literature searches using Scopus, PubMed and Web of Science (WoS) databases. The first one with the search terms “sexual conflict” & “male harm” OR “sexual conflict” & “female harm”, the second with the search terms “sexual conflict” & “female fitness” OR “sexual conflict” & “female productivity” OR “sexual conflict” & “female fecundity” OR “sexual conflict” & “female reproductive success”, and a final one with the search terms “sexual conflict” & “harassment”. Overall, we collected 121 effect sizes from 32 species and 51 studies.

We used the standardised mean difference with a small sample correction (Hedges’ *g* controlling for heteroscedastic population variances; hereafter SMDH) as our measure of effect size comparing female fitness across experimental treatments. We first fit multi-level meta-analytic models (intercept only) including study and species-level random effects. We also fit a model that included a species-level random effect with a phylogenetic correlation matrix, but the model containing only a species-level random effect variance was better supported, so we did not include phylogeny in our models. We applied robust variance estimators (RVEs) to correct standard errors from our models. Using our MLMA models, we also calculated effect size heterogeneity using *I*^2^. We explored drivers of effect size heterogeneity using multi-level meta-regression (MLMR) models, which included fixed effects (i.e., moderators) that we *a priori* predicted would impact female fitness: a) an index of sexual size dimorphism, b) the type of male harm, and c) sperm competition intensity, considering their interaction.

We chose the model with the lowest *AIC_c_* or the most parsimonious model. We fit all models with maximum likelihood for model comparison of fixed effect structure, and subsequently re-fit with restricted maximum likelihood when we identified the fixed effect structure. We explored publication bias using a new method that relies on fitting a MLMR model accounting for all the moderators available to explain variation in effects (random and fixed effects). More specific details about all analytical procedures can be found in the Supplementary Material.

### 2 Population genetic model

To test if an allele that confers males the ability to harm females can evolve independently of alleles that confer adaptation to the environment, we built a population genetic model considering a haploid population of females and males. All individuals carry two loci: an adaptation locus with two alleles (0 for the allele with an optimal phenotype to the environmental conditions, 1 for the allele with suboptimal phenotype to the environmental conditions), and a harm locus, expressed only on males, with two alleles (0 for the allele that makes individuals harm their sexual partners, 1 for the allele that makes individuals not harm their sexual partners). Individuals go through viability selection, where viability selection is stronger in individuals carrying the allele 1 than those carrying the allele 0. Surviving individuals become adults and enter the mating pool. Males with the allele 1 have higher mating success than males with the allele 0. During reproduction, males harm females in one of two different ways: a) mating harm, where harm is induced by the mating partner (e.g., traumatic insemination), and b) mating harassment, where harm to the females is induced by mating and non-mating males. After mating, a diploid zygote forms and recombination occurs between the two loci. Adults then die and new individuals are born.

Details about the derivation of the model can be found in the Supplementary Material. Briefly, assuming mating harassment, the change in frequency of the adaptation allele is

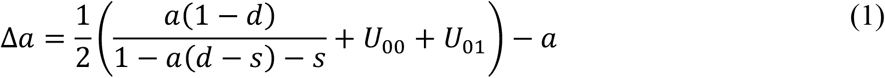

and the change in frequency of the harm allele is

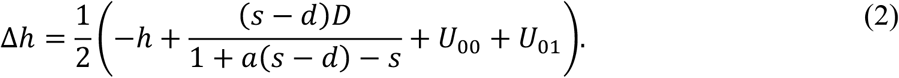

Assuming mating harm, the change in frequency of the adaptation allele is

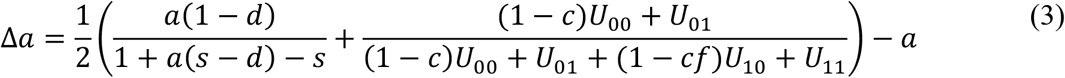

and the change in frequency of the harm allele is

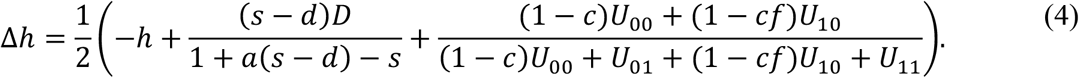

We analysed the model to find the evolutionary stable equilibrium of the population. In both cases, the change in linkage disequilibrium is Δ*D* = (*x*”_00_*x*”_11_ – *x*”_10_ *x*”_01_) – (*x*_00_ *x*_11_ –*x*_10_ *x*_01_), where *x*”_ij_ is the frequency of the genotypes in the next generation (with i, j = {0, 1}). Now we can solve the system of equations {Δ*a* = 0, Δ*h* = 0, Δ*D* = 0} to find the allele frequencies for which the population will no longer change. Unfortunately, Δ*D* is too complex to be solved and, therefore, we use a quasi-linkage equilibrium (Kimura, 1965) followed by a perturbation analysis to find approximated solutions to this system of equations. Finally, we do a stability analysis to find the stable equilibrium of the population (see SM).

## 3 Numerical simulations

We run a series of numerical simulations to track how male harm affects genetic adaptation and population recovery. To do this, we build a haploid genetic model of a population of males (*M*) and females (*F*) with the adaptation locus having two possible alleles: 0 for individuals adapted to the environment; and 1 for individuals not adapted to the environment. Importantly, from the results of the population genetic model, we know that the only evolutionary stable equilibrium is when both the adaptation allele and the harm allele are fixed in the population. Therefore, all males will be capable of harm but if there is an environmental change, only a minority will be adapted to the new environmental conditions. The numerical simulations follow the same life cycle as the population genetic model. With the numerical simulations we first tracked the population recovery and rate of genetic adaptation in both scenarios of sexual conflict, mating harassment and mating harm, when adapted males harm females to a higher degree than non-adapted males and when all males harm females at the same degree. Then, we explore the effect of sexual conflict in polygynous (i.e., males can mate with multiple females) and monogamous (i.e., males and females can only mate with one partner) populations. Detailed information regarding simulations can be found in the Supplementary Material. Briefly, under mating harassment, the birth rate of individuals with allele 0 and 1 is

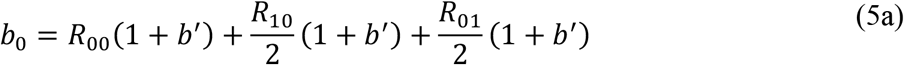

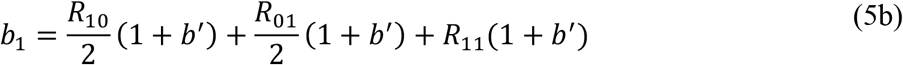

where *b*’ is the intrinsic birth rate, which is affected by the degree of male harm. We assume males with allele 0 impose higher costs than males with allele 1 by a scaling factor *f*.

Therefore, the strength of male harm depends on the frequency of the adapted allele in males,

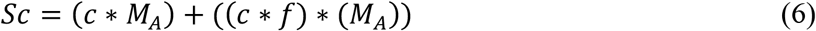

where *c* are the costs of sexual conflict imposed by males. Importantly, when *f* = 0 male harm is only imposed by adapted males, and when *f* = 1 adapted and maladapted males harm females to the same degree. Male harm is experienced equally by all females, regardless of their own allele, reducing female intrinsic birth rate (*b*’).

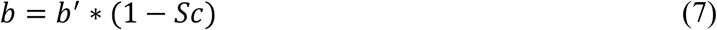

In the mating harm model, we assume that harm is not imposed via precopulatory mating harassment but during (e.g., traumatic insemination) or following copulation (e.g., transfer of harmful seminal proteins). Therefore, the costs of sexual conflict depend on the allele of the male mating with a female, and it is independent of the female allele. Then, the birth rate of adapted and maladapted alleles is

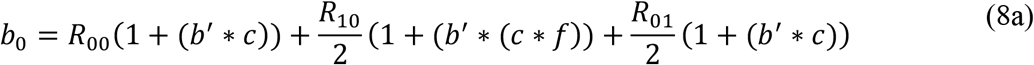

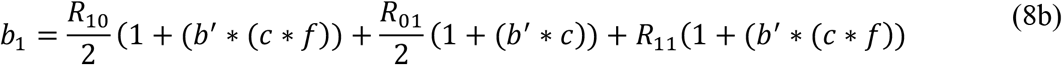

In all simulations presented here the population is initialized with a frequency of adapted alleles of 0.1. Moreover, we assume *ms* = 1.2, *d_A_* = 0.01, *d_a_* = 0.1 *and b_A_* = *b_a_* = 0.1. We run all simulations until populations recovered the initial population size (*N* = 100), or for 100 generations.

## Results

### Meta-analysis

Overall, we collected 149 effect sizes for a total of 32 species from 51 studies.

Unsurprisingly, invertebrates (classes: Insecta, Gastropoda, Malacostraca, Arachnida, Clitellata and Secernentea) made up most of the data (78.52%). We obtained 26 effect sizes from manipulations on species that resulted in direct harm (e.g., traumatic insemination), 60 from studies that manipulated indirect harm (e.g., mating harassment), and 63 effect sizes from experiments where females received both direct and indirect harm from male matings. We obtained 121 effects from 28 oviparous species, and 28 effects from 4 viviparous species.

Unfortunately, effect sizes from viviparous species were all taken from studies on fish with indirect male harm. As such, we analyzed only ‘harm type’ and an index of sexual size dimorphism.

#### Male harm negatively impacts female fitness

Experimentally manipulating female harm resulted in a strong decrease in female fitness overall (*i.e*., positive effect size with control group females having higher fitness than treatment groups: 0.59, 95% CI: 0.25 to 0.92). This effect held even when accounting for within-study non-independence (meta-analytic mean using Robust Variance Estimator: 0.59, 95% CI: 0.35 to 0.83), although publication bias suggest that results might be sensitive to additional studies (Supplementary Material). When accounting for sampling variance there was high effect size heterogeneity (*I^2^Total* = 0.91, 95% CI: 0.89 to 0.93) with most variance being the result of between study (*I^2^_study_* = 0.46, 95% CI: 0.34 to 0.57) and between species (*I^2^_Species_* = 0.23, 95% CI: 0.14 to 0.34) effects. Trait type and phylogeny explained much less variation overall (*I^2^_Trait_* = 0.17, 95% CI: 0.1 to 0.26; *I^2^_Phyiogeny_* = 0.05, 95% CI: 0.04 to 0.07; Table 1).

**Table 1:** Number of effect sizes, studies and species along with overall meta-analytic mean (average SMDH) and 95% confidence (CI) and prediction intervals (PI). Heterogeneity estimates and 95% CI’s are also provided for study, trait, species and total heterogeneity (excluding sampling variance)

| n<sub>effects</sub> | n<sub>studies</sub> | n<sub>species</sub> | Meta-analytic mean | 95% CI | 95% PI | I<sup>2</sup><sub>study</sub> | I<sup>2</sup><sub>trait</sub> | I<sup>2</sup><sub>species</sub> | I<sup>2</sup><sub>total</sub> |
| --- | --- | --- | --- | --- | --- | --- | --- | --- | --- |
| 149 | 51 | 32 | 0.59 | 0.35 to 0.83 | -1.28 to 2.45 | 45.59%<br>(33.88 - 57.33%) | 17.11%<br>(10.02 - 25.72%) | 22.96%<br>(13.73 - 33.85%) | 90.96%<br>(88.71 - 92.77%) |

#### Negative effects on fitness in relation to the type of male harm to females

Male harm type appeared to impact the overall magnitude of effects, explaining 12.12% of effect size variance (Fig. 1). Species with both direct and indirect harm showed a significantly higher impact on female fitness (meta-analytic mean: 0.85, 95% CI: 0.52 to 1.18, p = < 0.0001). Indirect harm resulted in a smaller but significant effect (meta-analytic mean: 0.51, 95% CI: 0.17 to 0.85, p < 0.01, Fig. 1). Surprisingly, species with only direct harm showed a small and opposite effect on female fitness (overall meta-analytic mean: −0.21, 95% CI: −0.81 to 0.4, p = 0.49; Fig. 1), but differences among species exhibiting different types of harm seemed to be driven primarily by a single study (Taylor *et al*., 2008) on *Drosophila simulans* (very high Cook’s distance). Removing this effect resulted in no significant difference between harm type categories.

**Figure 1.**
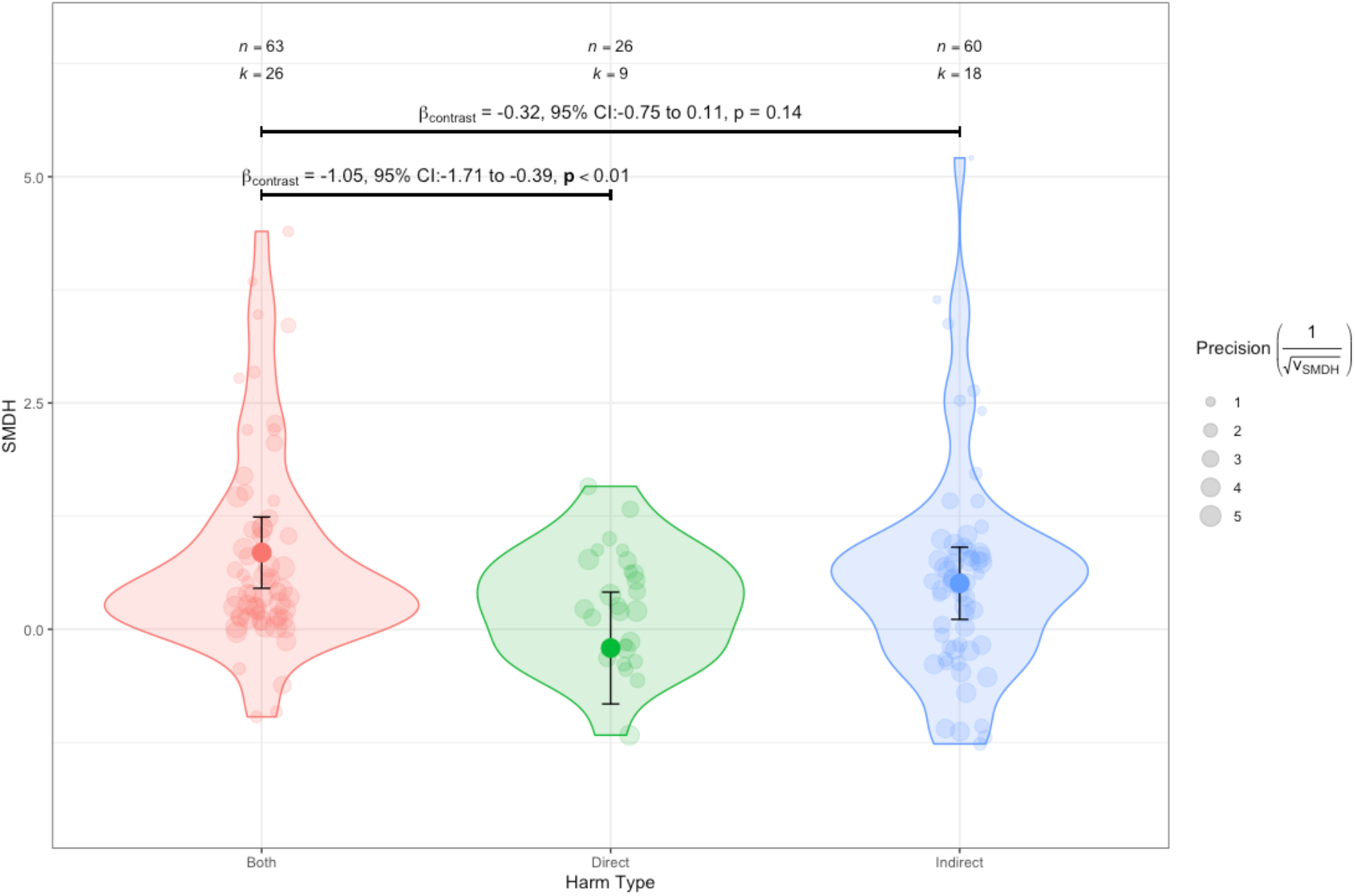
Overall, there we found a negative effect of male harm on female fitness, and this effect was dependent of the type of harm. Male harm to females was significatively different from 0 when harm was imposed indirectly (i.e., mating harassment) or by both direct and indirect harm, but not if the harm was imposed directly (e.g., traumatic insemination). Distribution of effect sizes, overall meta-analytic mean estimates and 95% confidence intervals (CIs) across species with different types of male harm towards females. Total effect sizes (n) and total number of studies (k) are provided for each level of male harm type. Relevant contrasts between meta-analytic means are provided, along with 95% CIs and significance of contrast. Note that this includes all data. Removing outlier point from single study results in no different between harm type categories.

#### Female fitness is more compromised with increasing SSD

As sexual size dimorphism (SSD) increases, female fitness is more negatively impacted by male harm (unstandardised slope, *β_SSD_* = 1.76, 95% CI = −0.05 to 3.56, p = 0.05, 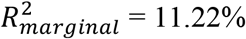; Fig. 2). In addition, there was no evidence that the effect of SSD varied according to the type of male harm exhibited (i.e., no interaction between Harm Type and SSD; *Δ_AIC_c__* =37.6, with the main effects model having the lowest *AIC_c_*). Interestingly, this pattern qualitatively appears to reverse in the one species where females are larger than males (*Idotea balthica*; Fig. 2).

**Figure 2.**
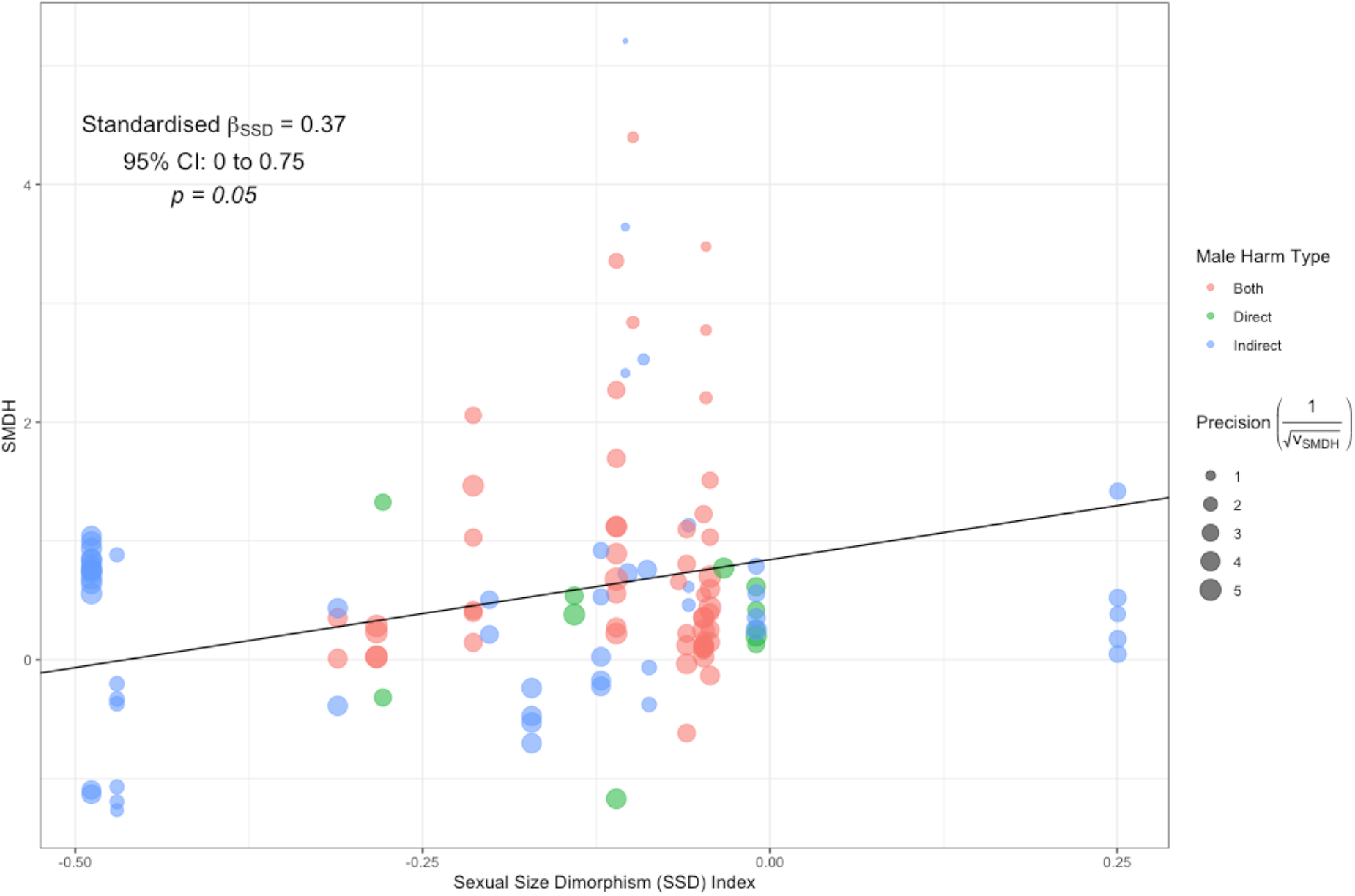
Female fitness is more strongly affected by male harm in species with large sexual size dimorphism. Note that although we provide an estimate of standardised slope (from a model with z-transformed SSD index) along with 95% CIs and significance in text, the regression line represents the slope of unstandardised SSD index along with raw data for ease of interpretation.

#### Male harm and sperm competition

Species with high sperm competition intensity exhibited a significant reduction in female fitness with male harm (meta-analytic mean: 0.68, 95% CI: 0.37 to 0.99, p = < 0.0001), whereas the same was not true for species with low sperm competition intensity (meta-analytic mean: 0.2, 95% CI: −0.19 to 0.6, p = 0.31) (Fig. 3). Overall, there was no significant difference between species deemed to have high sperm intensity compared to those with low sperm intensity.

**Figure 3.**
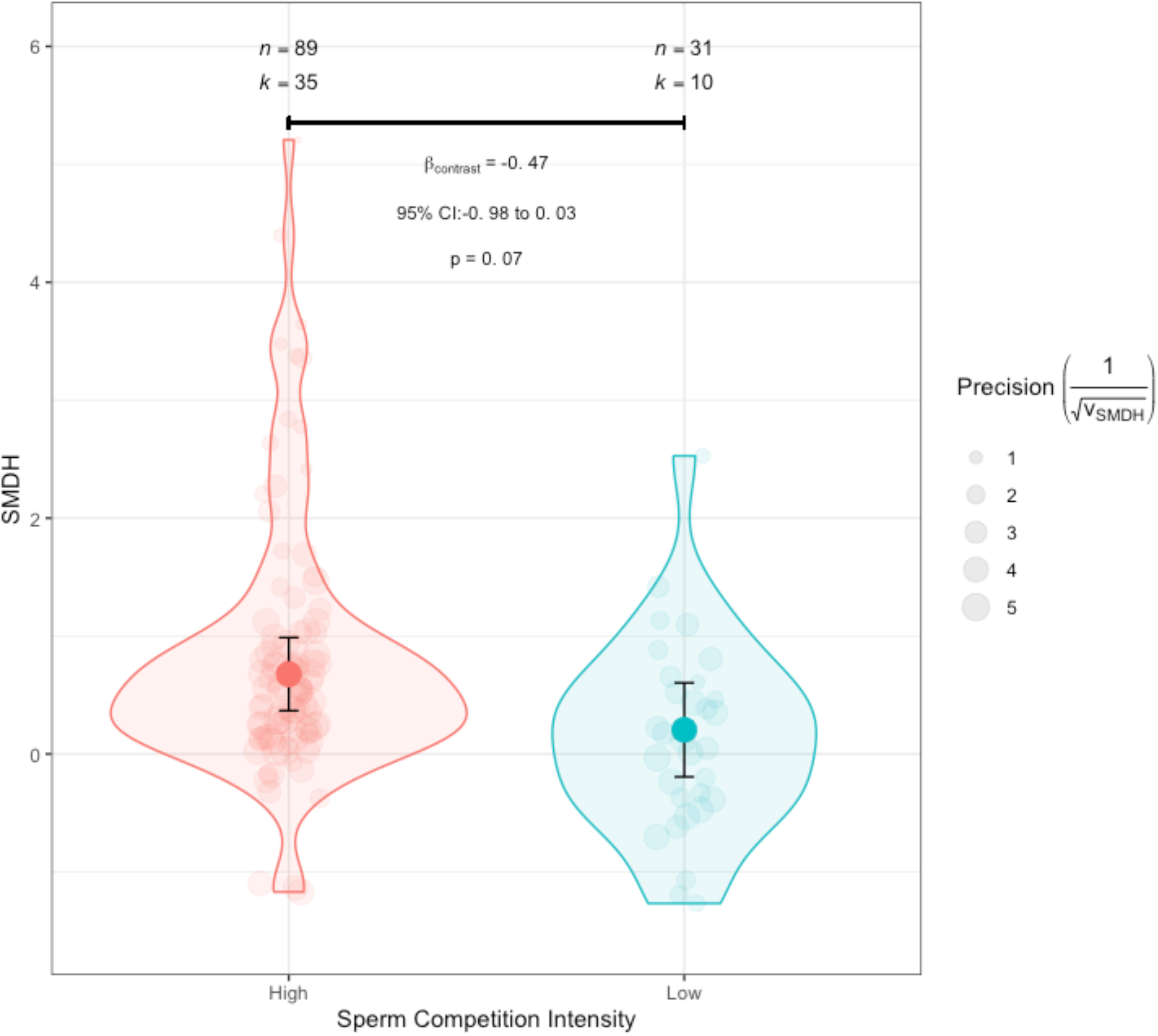
The effect of male harm on female fitness is dependent on sperm competition intensity. Overall, female productivity was affected in species with high sperm competition (p < 0.001) but the effect was not different from 0 in species with low sperm competition intensity. However, the difference between species with high and low sperm competition intensity was marginal.

### Population genetic model

Results from the population genetic model show that the only stable equilibrium for a population is when both the adaptation and harm alleles are fixed. This result is consistent across different levels of harm and the degree to which non-adapted males can harm the females, and is independent of whether harm comes from mating harassment or mating harm (Fig. 4). When non-adapted males are incapable of harming females *f* = 0; Fig. 4), the selective pressure for the harm allele to increase in the population is initially low. As the adaptation allele increases in the population, more males become capable of harming and, accordingly, of extracting the benefits of having the harm allele. At that point, the harm allele starts to increase in the population and the population only stops changing when both the adaptation and the harm alleles are fixed. Stability analysis confirms that this is the only stable point for this population.

**Figure 4.**
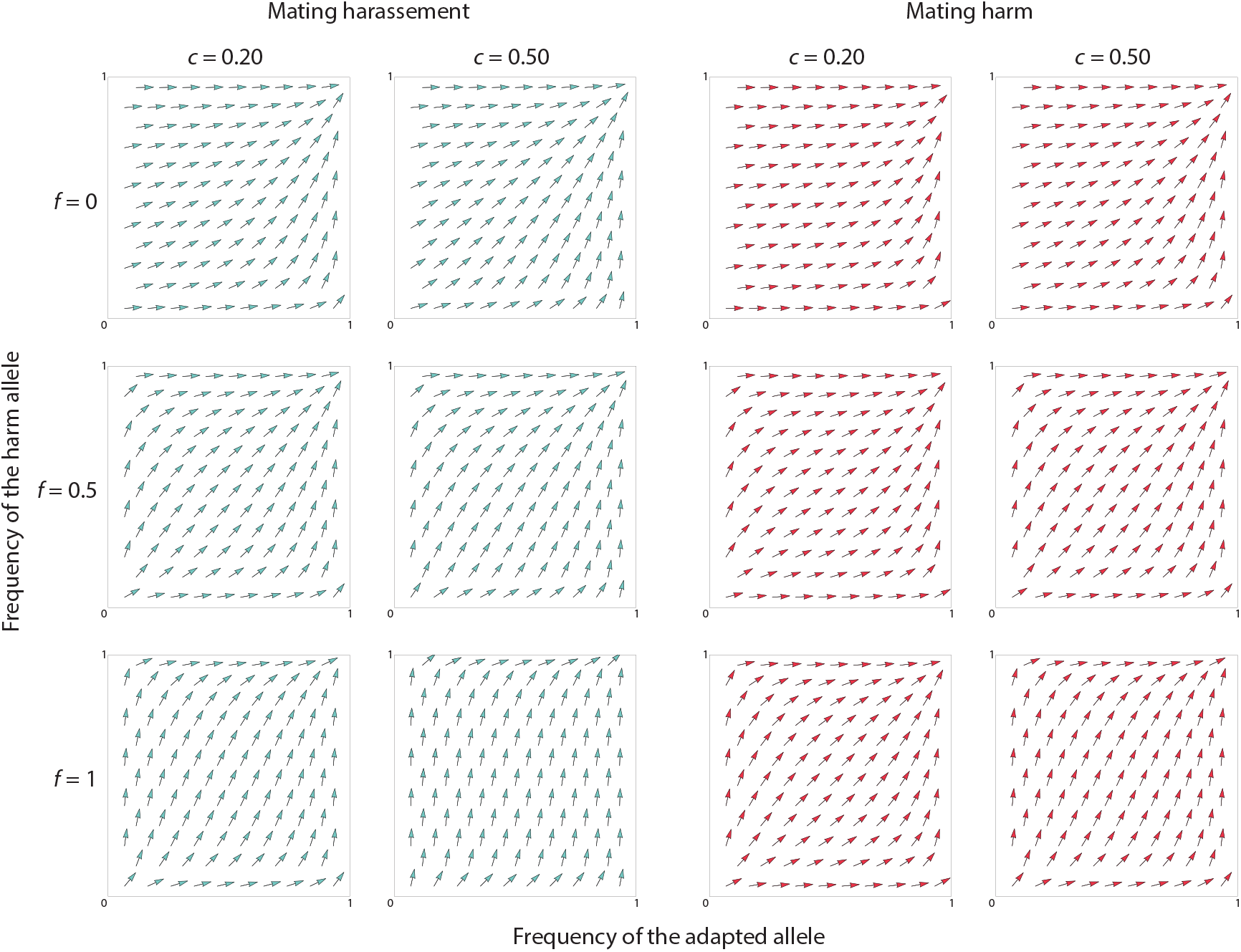
The only stable equilibrium is when both the adaptation allele and the harm allele are fixed in the population. Regardless of the cost of harm for females, the degree to which non-adapted males can harm the females, and the presence of mating harassment (right) or mating harm (left), the only stable equilibrium is when both the adaptation allele and the harm allele are fixed. Parameter values: viability cost of adapted males *d* = 0.01, viability cost of non-adapted males *s* = 0.10, recombination *r* = 0.5.

### Numerical simulations

#### Environmental effects on sexual conflict can facilitate evolutionary rescue by decreasing the impact on population demography

The mating harassment model shows that, if males with the adapted allele are more harmful than males with the non-adapted allele *f* < 1), population decline is reduced in the extent of the decline and the rate of the decline under both weak (*c* = 0.2) and strong sexual conflict (*c* = 0.5; Fig. 5). However, genetic adaptation is faster if all males impose equal costs *f* = 1). Interestingly, when sexual conflict is weak, populations recover faster if all males are equally harmful than if costs are dependent on the adapted allele, while there is no difference in the rate of recovery under strong sexual conflict (Fig. 5). The mating harm model shows some interesting differences with the mating harassment model. Namely, in the mating harm model, if males with the adapted allele are more harmful, the rate of population decline is reduced, but not the extent of the decline. Notably, when *f* < 1, because populations decline more slowly and maladapted alleles can remain for longer, genetic adaptation and population recovery is slower under both weak (*c* = 0.2) and, more markedly, strong sexual conflict (*c* = 0.5; Fig. 5). Therefore, the effects of sexual conflict on evolutionary rescue not only depend on the strength but also on the mechanism of sexual conflict.

**Figure 5.**
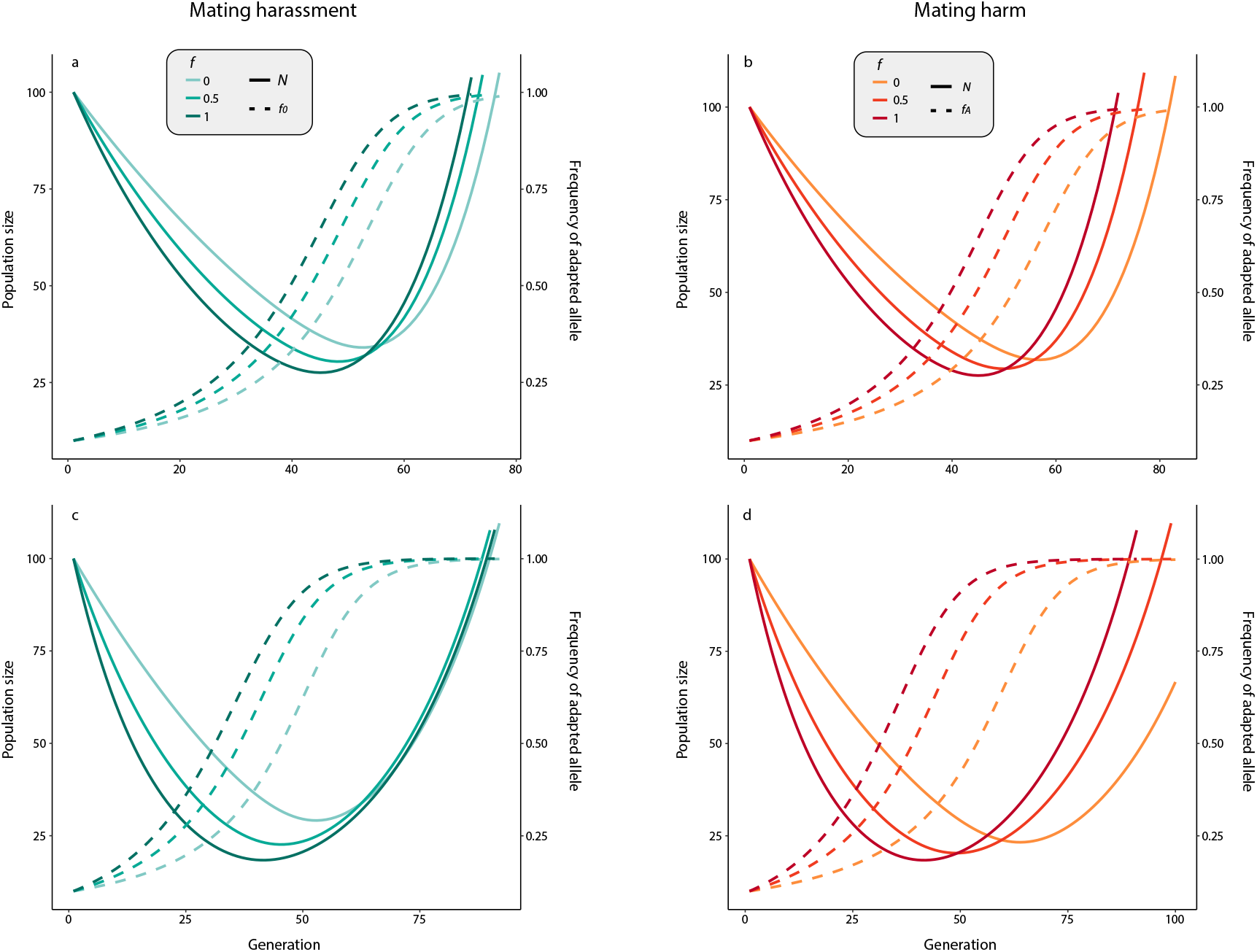
The effect of sexual conflict on evolutionary rescue depends on the strength and mechanism by which the costs are imposed on females. Mating harassment reduces population decline, but slows adaptation, if it is higher in adapted vs. non-adapted males (**A, C**). Importantly, when sexual conflict is weak (*c* = 0.2; **A**), populations recover faster if the level of male harm is similar between adapted and non-adapted males, but there is no difference if sexual conflict is strong (*c* = 0.5; **C**). When the costs of sexual conflict are imposed trough mating harm (**B, D**), higher harm by adapted males reduces the rate of population decline, but not the extent of this decline. However, populations take longer to recover if the costs are imposed more strongly by adapted males under both low (*c* = 0.2; **B**) and high sexual conflict (*c* = 0.5; **D**), albeit more clearly in the latter case. Note that x axes differ between the top and bottom row. This is because, under weak sexual conflict, populations recover their initial size much faster than under strong sexual conflict, especially in the case of mating harm. The solid lines reflect population size (*N*) and the dashed line depicts the frequency of the adapted allele (0). The scaling factor *f*) reflects the costs of sexual conflict imposed by males with the non-adapted allele relative to males with the adapted allele *f* = 1 all males impose equal sexual conflict costs, *f* = 0.5 males with the non-adapted allele impose half the costs than males with the adapted allele, and *f* = 0 only males with the adapted allele impose sexual conflict costs). In these simulations, *h* = 0, representing extreme polygyny.

#### The demographic benefits of sexual conflict decrease in less polygynous populations

In the mating harassment model, polygynous populations decline less than less polygynous populations (Fig. 6). Polygyny also facilitated population recovery; population size after 100 generations was twice as large in polygynous over monogamous populations (Fig. 6). The rate and extent of population decline was minimized when sexual conflict was imposed only by adapted males (*f* = 0), although this effect was small. We found a similar pattern in the mating harm model, except that in this case only the rate, and not the extent, of population decline was minimized (Fig. 6). There were no large differences in the rate of genetic adaptation between polygynous and monogamous populations in either the mating harassment or mating harm model (Fig. 6).

**Figure 6.**
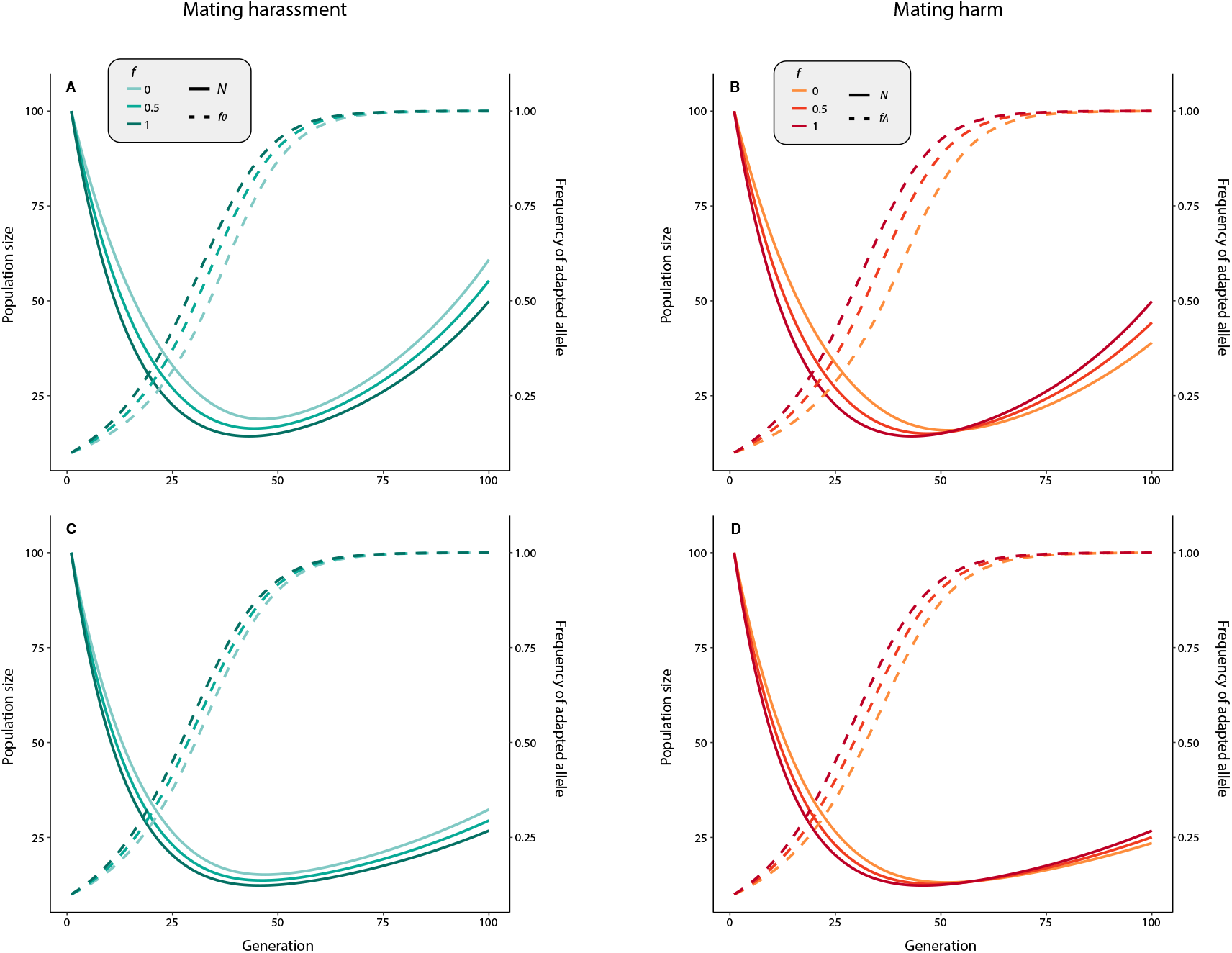
Sexual conflict is less likely to aid evolutionary rescue in less polygynous populations. In the mating harassment model, more polygynous populations recover faster (*h* = 0.5, **A**) than monogamous populations (*h* = 1, **C**), but the benefit of environmental effects on sexual conflict are small. In the mating harm model, although polygynous populations recovered faster (*h* = 0.5, **B**) than monogamous populations (*h* = 1, **D**), the effect of environmental effects on sexual conflict was greatly reduced. In all simulations sexual conflict is weak (*c* = 0.2). Shown are population size (*N*) in solid line and the frequency of the adapted allele (0) in broken line.

## Discussion

Whether species can adapt fast enough to prevent extinction is a pressing question, especially given the rates of anthropogenic climate change. Although sexual selection has been shown to aid rapid adaptation, it is often associated with sexual conflict, which can have detrimental consequences for female and population fitness, and facilitate extinction. We provide evidence that sexual conflict not only has a negative effect on female fitness across taxa, but can affect the evolutionary dynamics of populations, reducing population decline at the cost of slowing the process of adaptation. Therefore, our study suggest that sexual conflict can be an important factor affecting the response of species to climate change.

Our meta-analysis on available studies that have manipulated male harm levels and studied associated female fitness shows that male harm decreases female fitness across taxa. While evidence to this respect has been accumulating in some species for the last few decades (Parker, 1979; Arnqvist & Rowe, 2005), several studies have failed to find a net impact of male harm on female fitness in some species (*e.g*., Mouginot *et al*., 2015; Nakata, 2016). Our findings suggest that male harm indeed generally negatively effects on female productivity across studied taxa. Interestingly, our results suggest that mating harm (i.e., indirect harm) may be more costly for females. However, the lack of definitive differences among harm type categories might actually be expected given that male sexual traits that are considered harmful for females, such as genital spines, can have beneficial effects in some species by increasing female productivity (*e.g*., Arnqvist *et al*., 2021). Our results also suggest that the net effect of male harm on female fitness may be affected by sexual size dimorphism and sperm competition. Although results are preliminary given the relative scarcity of data to this respect, this suggests an association between stronger sexual selection and higher male harm to females, which is in accordance with theoretical expectations.

Establishing that male harm is deleterious for female fitness raises the intriguing question as to how this may affect population adaptability to novel environments, and how male harm generally interacts with the process of adaptation. We show how the interaction between sexual selection and sexual conflict has the potential to affect the dynamics of evolutionary rescue of a population. Our models suggest that sexual conflict can limit population decline, but at the cost of slower adaptation. Our results, therefore, suggest that sexual conflict can act like a double edge sword, reducing extinction risk by buffering population decline but delaying genetic adaptation. Given that sexual conflict reduces female fecundity, it is not surprising that, under strong sexual conflict, population decline is steeper than under weak sexual conflict. Thus, when sexual conflict depends on male adaptation to the environment, the overall impact on female fecundity (and hence population decline) is reduced, in agreement with empirical evidence (García-Roa *et al*., 2019). Our models further show that reduced sexual conflict can maintain larger populations in which maladapted alleles can remain for longer, slowing down the process of adaptation. This, however, is only true if sexual conflict occurs through mating harassment. Polygynous populations exhibiting mating harassment might be therefore better equipped to adapt and persist in the face of climate change. Importantly, our model assumes a sudden or rapid environmental change, and evolutionary rescue is more likely under gradual change (Bell & Gonzalez, 2011; Carlson *et al*., 2014), although faster recovery might be at the cost of long-term survival (Liukkonen *et al*., 2021). Exploring the effects of sexual conflict in gradually changing environments would be an interesting expansion.

A major assumption of our model is that sexual conflict is imposed more strongly by adapted males, an assumption that has been used theoretically (Bonduriansky, 2014; Connallon, 2015; Connallon & Hall, 2018) and supported empirically (Long *et al*., 2012; Chenoweth *et al*., 2015; García-Roa *et al*., 2019). Based on this evidence, we believe our model reflects realistic biological scenarios, although the magnitude and generality of this assumption still needs further investigation. It is worth noting that we do not take into account female resistance, which reduces the costs of male harm (Rice, 1996; Holland & Rice, 1999). However, irrespective of whether females can evolve resistance attenuating the impact of sexual conflict, the relevance of our model relies in the difference in the costs imposed by adapted and non-adapted males independent of female resistance. What is less known is whether resistance differs in more and less adapted females, which could in principle impact the dynamics of evolutionary rescue and should be subject to experimental scrutiny.

An interesting expansion to our models is the case of assortative mating. Previous studies have shown that sexual conflict can be directed towards high fecundity females (i.e., better adapted) (Long *et al*., 2009; Chenoweth *et al*., 2015), and that males in good condition (i.e., better adapted) can prevent males in poor condition access to females (Gómez-Llano *et al*., 2020). In such case, the costs of sexual conflict would act only in a subset of the population (i.e., adapted females), which could reduce the variance in female fitness and the efficiency of selection eliminating maladapted alleles. Similarly, kin selection may reduce the harm that males impose on females (Rankin, 2011; Carazo *et al*., 2014; Faria *et al*., 2015, 2020), reducing the fitness difference between adapted and non-adapted males. Therefore, while kin selection has been implied in reducing population costs from sexual conflict, it could maintain non-adapted alleles for longer periods in a population.

Given the current rate of human-induced environmental change and its potential effects in decreasing population size and increasing extinction risk across species (Dirzo *et al*., 2014; Ceballos *et al*., 2017; Wagner *et al*., 2021), understanding what mechanisms modulate the processes of adaptation and population recovery is a pressing need in evolutionary and conservation biology. We have shown here that sexual conflict can be important to understand evolutionary rescue and population extinction at large. Our study underscores the need for a better understanding of the ecology of sexual conflict and its consequences for adaptive processes. Our models are a necessary but preliminary step in a direction were clearly more theoretical and empirical research is necessary. Ultimately, some degree of biodiversity loss in response to climate change is inevitable, but mitigation of this loss will require an efficient use of conservation resources. To do that, it is vital to understand the processes that better equip species to adapt. Our work generates new sets of hypotheses that, we hope, may further both theoretical and empirical research.

## Acknowledgements

MGL was supported by the NSF (DEB 1748945). G.S.F was funded by a Leverhulme Trust Early Career Fellowship. PC was supported by grant PID2020-118027GB-I00 funded by MCIN/AEI/10.13039/501100011033 and by a 2018 Leonardo Grant for Researchers and Cultural Creators, BBVA Foundation.

## Data accessibility

All data will be available in a public repository upon acceptance

