## Supplementary Material for "Male harm suppresses female fitness to affect the dynamics of adaptation and evolutionary rescue"

**Supplementary Material: Male harm supresses female fitness to affect the dynamics of adaptation and evolutionary rescue**

Miguel Gómez-Llano, Gonçalo S. Faria, Roberto García-Roa, Daniel W.A. Noble, Pau Carazo

**Methods**

***1- Meta-analysis***

#### *1.1- Systematic literature search*

We conducted a systematic review of the existing literature following the PRISMA protocol (Liberati *et al.*, 2009) as closely as possible. Specifically, we looked for studies that experimentally manipulated the level of male harm to females and measured its outcome in terms of female fitness (i.e., described adaptations leading to male harm to females, consisting of male adaptations involving direct trauma to females. We only qualified extracted phenotypic traits when it was clear from the reported paper, or the raw data, that the trait had a direct negative impact on female lifetime reproductive success and/or (in the absence of this measures) because male adaptations inflicted obvious injuries to females. Due to the co-evolution of female resistance and male harm, harmful male adaptations may not be expected to impose high fitness costs in females over most evolutionary time (Reinhardt *et al.*, 2015). We thus opted to include both cases where the consequences of male harm were measured in terms of female fitness (i.e. quantitative evidence) and cases in which lifetime/reproductive fitness costs to females were not studied but male adaptations involved produced measurable harm to females (i.e. injuries), such as in traumatic insemination via genital ablation or copulatory wounding, or in cases where male harassment regularly leads to female injuries and occasional deaths (i.e. qualitative evidence). We conducted a first literature search on 03/04/20 using the Scopus, PubMed and Web of Science (WoS) databases with the search terms “sexual conflict” & “male harm” OR “sexual conflict” & “female harm” for animal taxa. Overall, very few papers were found with these search strings (73 total: Scopus = 31, PubMed = 15 and WoS = 27). After removing duplicates only 36 papers were relevant, and we exported them to Rayyan. We conducted a second literature search on 03/04/20 using the Scopus, PubMed and Web of Science (WoS) databases with the search terms: “sexual conflict” & “female fitness” OR “sexual conflict” & “female productivity” OR “sexual conflict” & “female fecundity” OR “sexual conflict” & “female reproductive success”. We found a total of 694 papers (Scopus = 250, PubMed = 144 and WoS = 300). After removing 373 duplicates, we exported 321 to Rayyan. We conducted a final search on the 7/04/20 using the search terms: “sexual conflict” & harassment. We found a total of 414 items (Scopus = 175, PubMed = 50 and WoS = 189). After removing 175 duplicates, we exported 239 to Rayyan. In Rayyan, we checked for duplicates within the complete database comprising all the papers located via these three searches and removed 69 duplicates, leading to 527 unique studies for more detailed screening. Based on the title and abstract we excluded 347 papers that clearly did not report adaptations for male harm, leaving a total of 180 papers for in-depth screening. We carefully screened these papers and excluded papers that did not comply with our selection criteria described above. In the process of screening, we added 27 more papers through forward and backward searches of citations and references, leading to a total final sample of selected studies reporting male harm adaptations for a total of 87 different species (see supplementary materials for complete list).

Overall, we collected effect sizes for a total of 32 species from 51 studies. Unsurprisingly, invertebrates (classes: Insecta, Gastropoda, Malacostraca, Arachnida, Clitellata and Secernentea) made up most of the data (78.52%). We obtained 26 effect sizes from manipulations on species that resulted in direct harm (e.g., traumatic insemination), whereas 60 effects came from studies on species that manipulated indirect harm to females (e.g., mating rate and harassment). A total of 63 effect sizes were from experiments where females received both direct and indirect harm from male matings. Across all species, we obtained 121 effects from 28 oviparous species, and 28 effects from 4 viviparous species. Unfortunately, effect sizes from viviparous species were all taken from studies on fish species with indirect male harm. As such, we analyzed only ‘harm type’ and an index of sexual size dimorphism.

#### *1.2- Standardised Effect Size Measure*

We used the standardised mean difference with a small sample correction (i.e., Hedges’ *g*) as our measure of effect size comparing female fitness measures across experimental treatments. To correct for the possibility that population variances in the two treatments differ, we made use of a corrected version of Hedges’ *g* that controls for heteroscedastic population variances (hereafter called SMDH). Effect sizes were calculated by subtracting the control group mean from the treatment group mean and dividing by the pooled standard deviation. As such, positive effect sizes indicate that female ‘fitness/traits’ in control groups was higher than female ‘fitness/traits’ in treatment groups.

#### *1.3- Meta-analysis*

We analysed effect size data using the *metafor* (Viechtbauer, 2010) package in R (R Development Core Team, 2018) (vers 4.0.5). To estimate the overall effect of male-harm on female fitness we first fit multi-level meta-analytic models (intercept only). We plotted overall meta-analytic means using the *orchaRd* package (Nakagawa *et al.*, 2021). Overall, we collected between 1 to 12 effect sizes per study. To control for sources of non-independence, we included study and species-level random effects. We also fit a model that included a species-level random effect with a phylogenetic correlation matrix derived using the Open Tree of Life Database (https://tree.opentreeoflife.org/). We used Grafen’s method (Grafen, 1989) to calculate branch lengths using the R package *ape* (Paradis & Schliep, 2019). Overall, a model containing only a species-level random effect variance was better supported than a model that estimated a phylogenetic variance ($\Delta_{AIC_{c}}$ = 2.16). As such, we did not include phylogeny in our models.

Study and species-level random effects cannot account for additional sources of within-study non-independence, such as effect sizes calculated using common treatment groups or for different traits measured on the same individuals (Noble *et al.*, 2017). We therefore applied robust variance estimators (RVEs) to correct standard errors from our models. RVEs have been shown to be excellent estimators of standard errors from meta-analytic models when effect sizes within studies are correlated. In addition, one does not need to assume, or specify, how such effects are correlated, making them easier to apply.

Using our MLMA models we also calculated effect size heterogeneity using $I^{2}$. $I_{study}^{2}$ and $I_{species}^{2}$ are defined as the proportion of effect size variance over total variance as a result of between study and species effects, respectively. In contrast, $I_{total}^{2}$ was calculated as the proportion of total effect size variance after accounting for, or removing, total sampling variance.

We explored drivers of effect size heterogeneity using multi-level meta-regression (MLMR) models. Our models included fixed effects (i.e., moderators) that we *a priori* predicted would impact female fitness. More specifically, we predicted that the intensity of sexual selection and sexual conflict would affect how much harm males did to females. To test these predictions, we included an index of sexual size dimorphism (SSD), which involved taking the log transformed ratio between male body size to female body size for the species (i.e., $log\left( \frac{Male_{bodysize}}{Female_{bodysize}} \right)$, hereafter referred to as SSD). Positive SSD values indicate species with larger males compared to females, whereas negative SSD values indicate species where females are larger than males. While this does not directly measure the intensity of sexual selection, it is a proxy for it, and has been used in meta-analyses in the past (*e.g.,* Janicke & Fromonteil, 2021).

In addition to SSD, we also categorised effect sizes as having come from species with different types of male harm and sperm competition intensity (see moderators section). These included species with direct male harm (i.e., species with traumatic insemination) as well as species with indirect male harm (i.e., species that harm females through harassment). There are also a number of species where it is clear both direct and indirect forms of male harm exist. We also fit separate models to a two-level categorical moderator describing the sperm competition intensity of the species (i.e, “high” or “low”). In both these models, we controlled for SSD index, but included z-transformed SSD, such that meta-analytic means in each categorical moderator are for species of average SSD. We also accounted for the possibility that SSD might vary depending on the specific harm type and sperm competition intensity (i.e., interaction between SSD and Harm type and Sperm competition intensity). We evaluated support for interactions by comparing a model including the interaction to a model without it, comparing $AIC_{c}$ between the two models. We chose the model with the lowest $AIC_{c}$ if they differed by greater than 2, otherwise, we choose the most parsimonious model. All models comparing fixed effects were fit with maximum likelihood for model comparison of fixed effect structure, and then subsequently re-fit with restricted maximum likelihood when the fixed effect structure was identified.

Given that MLMR models assume homogeneity of variances across the levels of categorical moderators, and this assumption appeared to be violated when visually inspecting the data, we also fit a model that assumed heterogeneous residual variance across the different levels of male harm type moderator (i.e., different residual variance for ‘Both’, ‘Direct’ and ‘Indirect’ male harm type). We compared this model to a homogeneous variance model using $AIC_{c}$. Overall, the heterogeneous variance models did not improve overall fit (Harm Type: $\Delta_{AIC_{c}}$ = 0.63; Sperm Competition Intensity: $\Delta_{AIC_{c}}$ = 0.72), and so, we report results from models assuming homogeneity of variance.

#### *1.4 Publication Bias*

Publication bias results when studies with lower statistical power that find opposite patterns to the predicted effects (i.e., male harm increases female fitness) go unpublished. This can result in effects being upwardly biased overall. Evidence for publication bias can be roughly detected by inspecting a funnel plot for funnel asymmetry. However, any asymmetry identified may simply be the result of high effect size heterogeneity. As such, to formally test for publication bias we used a new method that relies on fitting a MLMR model accounting for all the moderators available to explain variation in effects (i.e., all random and fixed effects) (Nakagawa *et al.*, 2022). Using this model, we also include effect size sampling variance as a moderator. This approach has the benefit of being a formal way to both statistically test for publication bias (while accounting for as much effect size heterogeneity as possible) and correcting for it; providing a sensitivity analysis on how the effect is expected to change if we were to hypothetically observe the missing effects (Nakagawa *et al.*, 2022). It is still important to note that this corrected overall mean should be interpreted with caution. We can never know exactly how many studies, if any, are missing. As such, these should be viewed as a sensitivity analysis.

***2- Population genetic model***

*2.1- The model.*

We consider a haploid population of females and males. All individuals carry two loci: an adaptation locus with two alleles (0 for the allele with an optimal phenotype to the environmental conditions, 1 for the allele with suboptimal phenotype to the environmental conditions), and a harm locus with two alleles (0 for the allele that makes individuals harm their sexual partners, 1 for the allele that makes individuals not harm their sexual partners). Importantly, the harm locus is only expressed by males. Accordingly, the frequency of the adaptation allele is *a* = *x*_00_ + *x*_01_ and the frequency of the harm allele is *h* = *x*_00_ + *x*_10_, where *x*_ij_ is the frequency of the genotype in the population (with i, j = {0, 1}). The linkage disequilibrium between the two loci is *D* = *x*_00_ *x*_11_ – *x*_10_ *x*_01_.

Individuals carrying allele 0 for the adaptation locus suffer viability cost of *d* while individuals carrying allele 1 suffer a viability cost of *s* (with *d* < *s*). Accordingly, genotype frequencies change to *x*’_00_ = (*a h* + *D*) (1 – *d*)/(*a*(1 – *d*) + (1 – *a*)(1 – *s*)), *x*’_01_ = (*a* (1 – *b*) – *D*) (1 – *d*)/(*a*(1 – *d*) + (1 – *a*)(1 – *s*)), *x*’_10_ = ((1 – *a*) *h* – *D*) (1 – *s*)/(*a*(1 – *d*) + (1 – *a*)(1 – *s*)), and *x*’_11_ = ((1 – *a*) (1 – *h*) + *D*) (1 – *s*)/(*a*(1 – *d*) + (1 – *a*)(1 – *s*)), where *a*(1 – *d*) + (1 – *a*)(1 – *s*) is the average viability in the population. After viability selection, males harm females. Harm occurs in one of two different ways: a) mating harm, where harm to the females is induced during (e.g., traumatic insemination) or immediately after (e.g., toxic ejaculates) mating, and b) mating harassment, where harm to the females is induced before mating by mating and non-mating males.

Harmful males, the ones carrying allele 0 for the harm locus, always have an advantage over non-harmful males. However, how successful harmful males are depends on whether they carry allele 0 for the adaptation locus. Males that carry allele 0 for the adaptation locus can enjoy the full benefits of their harm while males that carry allele 1 only enjoy a proportion *f* of the benefits of their harm (meaning that they also only harm the females a proportion *f* of the harm that they would be able to inflict otherwise). If it is mating harm, only females mating with harmful males receive the cost *c* of harm, while if it is mating harassment all females receive the cost *c* of harm, regardless of their mating partner. The mating success of the different types of males in the population is therefore *U*_00_ = *k* *x*’_00_ / (*k* *x*’_00_ + *x*’_01_ + *m* *x*’_10_ + *x*’_11_), *U*_01_ = *x*’_01_ / (*k* *x*’_00_ + *x*’_01_ + *m* *x*’_10_ + *x*’_11_), *U*_10_ = *m* *x*’_10_ / (*k* *x*’_00_ + *x*’_01_ + *m* *x*’_10_ + *x*’_11_), and *U*_11_ = *x*’_11_ / (*k* *x*’_00_ + *x*’_01_ + *m* *x*’_10_ + *x*’_11_), where *k* = (1 + *c*) / (1 – *c*) and *m* = (1 + *c f*) / (1 – *c f*). Accordingly, the more harmful males are, the more mating success they have. After mating, there is a formation of a diploid zygote, and recombination occurs between the two loci with probability *r*. Adults then die and new individuals are born.

We now calculate how the allele frequencies change from one generation to the next. Assuming mating harassment, the change in frequency of the adaptation allele is

$$\Delta a=\frac{1}{2}\left( \frac{a\left( 1-d \right)}{1-a\left( d-s \right)-s}+U_{00}+U_{01} \right)-a$$

(1)

and the change in frequency of the harm allele is

$$\Delta h=\frac{1}{2}\left( -h+\frac{\left( s-d \right)D}{1+a\left( s-d \right)-s}+U_{00}+U_{01} \right).$$

(2)

Assuming mating harm, the change in frequency of the adaptation allele is

$$\Delta a=\frac{1}{2}\left( \frac{a\left( 1-d \right)}{1+a\left( s-d \right)-s}+\frac{(1-c)U_{00}+U_{01}}{\left( 1-c \right)U_{00}+U_{01}+(1-cf)U_{10}+U_{11}} \right)-a$$

(3)

and the change in frequency of the harm allele is

$$\Delta h=\frac{1}{2}\left( -h+\frac{\left( s-d \right)D}{1+a\left( s-d \right)-s}+\frac{(1-c)U_{00}+(1-cf)U_{10}}{\left( 1-c \right)U_{00}+U_{01}+(1-cf)U_{10}+U_{11}} \right).$$

(4)

In both cases, the change in linkage disequilibrium is Δ*D* = (*x’’*_00_ *x’’*_11_ – *x’’*_10_ *x’’*_01_) – (*x*_00_ *x*_11_ – *x*_10_ *x*_01_), where *x’’*_ij_ is the frequency of the genotypes in the next generation (with i, j = {0, 1}). Now we can solve the system of equations {Δ*a* = 0, Δ*h* = 0, Δ*D* = 0} to find the allele frequencies for which the population will no longer change. Unfortunately, Δ*D* is too complex to be solved and, therefore, we use a quasi-linkage equilibrium (Kimura, 1965) followed by a perturbation analysis (see 2.2 below) to find approximated solutions to this system of equations. Finally, we do a stability analysis to find the stable equilibrium of the population (see 2.3 below).

*2.2- Quasi-linkage equilibrium*

When linkage disequilibrium is too complicated to be solved using a standard approach, we can get an approximation by using quasi-linkage equilibrium (QLE)(Kimura, 1965). Using QLE allows us to settle the value of linkage disequilibrium (*D*) to an approximated constant value (*D*_Q_) provided that the dynamics that occur through natural selection take place at a slower rate than the changes in the value of linkage disequilibrium through recombination. To find the QLE value of *D*, the condition *D’’* = *D* = *D*_Q_ needs to be solved, where *D’’* is the value of linkage disequilibrium following selection, recombination, and mutation. This condition cannot be solved explicitly, but we can get an approximation for *D*_Q_ by using a perturbation analysis (an accessible account is provided in(Otto & Day, 2007), Ch. 9). To make explicit that selection is weak, we multiply every selection coefficient by a small value $\delta$. The QLE condition now becomes *f*($\delta$) = *D*_Q_, where *D*_Q_ = *D*_0_ + *D*_1_ $\delta$ + *D*_2_ $\delta$^2^ + … and solve the equation *f*($\delta$) – *D*_Q_ = 0. By using the Taylor series of *f*($\delta$) to first order in $\delta$ near the point $\delta$ = 0, we get

$$f\left( \delta\right)=f\left( 0 \right)+\left( \frac{df}{d\delta}|_{\delta=0} \right)\delta=0$$

(5)

and the terms in the approximation for *D*_Q_ can be found by solving each term in the (M1) to zero and solving for *D*_i_. The following *D*_Q_ is obtained:

$$D_{Q}=\frac{\left( 1-a \right)a\left( 1-b \right)b c(1-f)(1-r)}{2r}$$

(6)

where *r* is the recombination value.

*2.3- Stability analysis*

To determine the stability of an equilibrium point, we first calculate the Jacobian matrix **J** (Otto & Day, 2007). Each entry of the matrix is given by:

$$J=\frac{\partial p}{\partial q}|_{p,q=0}$$

(7)

where *p* = {$\Delta$*a*,$\Delta$*h*,$\Delta$*D*} and *q* = {*a*,*h*,*D*}. If the leading eigenvalue of this matrix is negative, then the point is considered stable.

***3- Numerical simulations***

We build a haploid genetic model of a population of males (*M*) and females (*F*) with the adaptation locus having two possible alleles: 0 for individuals adapted to the environment; and 1 for individuals not adapted to the environment. Importantly, from the results of the population genetic model, we know that the only evolutionary stable equilibrium is when both the adaptation allele and the harm allele are fixed in the population. Therefore, all males will be capable of harm but if there is an environmental change, only a minority will be adapted to the new environmental conditions.

First, we tracked population recovery and adaptation in both scenarios of sexual conflict, mating harassment and mating harm, when adapted males harm females to a higher degree than non-adapted males and when all males harm females at the same degree. Then, we explore the effect of sexual conflict in polygynous (i.e., males can mate with multiple females) and monogamous (i.e., males and females can only mate with one partner) populations.

The growth rate of the population is given by

$$g_{z}=(1+b_{z})(1-d_{z})$$

(8)

where $b_{z}$ is the birth rate and $d_{z}$ the intrinsic death rate of individuals with allele *z*, where $z=\left\{ 0, 1 \right\},$ $b_{0}=b_{1}$ and $d_{0}<d_{1}$.

The adapted allele also gives males higher mating success by increasing the frequency of the adapted male alleles that go into mating by a factor *m* (being *m >* 1). The frequency of adapted alleles in males is then

$$M_{0}^{'}=\frac{M_{0}*m}{{(M}_{0}*m)+(1-M_{0})}$$

(9)

Mating is determined by the frequencies of the different genotypes in the population, and the frequency of matings is given by

$$R_{z,w}= \frac{M_{z}F_{w}}{\sum_{z}^{w} (M_{z}^{'}*{(F}_{w}*h))}$$

(10)

where *z*, *w* = {0, 1}. Here, *h* controls the number of mating pairs a male can have, allowing us to test the role of mating system, different levels of polygyny and monogamy, in evolutionary rescue. Specifically, 0 <= *h* <= 1 being *h* =1 monogamy and *h* = 0 extreme polygyny. Sexual conflict affects female fecundity through mating harassment or mating harm (see main text for equations).

**Results**

**Meta-analysis: estimating biases**

Visual inspection of funnel plot asymmetry suggested evidence for publication bias (i.e., missed effects sizes with low precision when female fitness was worse in control treatments) (Figure S1), which was confirmed through the identification of a significant slope between effect size and effective sample size (*β* = 2.13, 95% CI: 0.56 to 3.71). This result held true when accounting for heterogeneity using meta-regression models (*β* =2.27, 95% CI: 0.56 to 3.98). Correcting for the possibility of missing studies resulted in the overall meta-analytic mean effect size being indistinguishable from zero (Corrected meta-analytic mean: -0.14, 95% CI: -0.78 to 0.5).

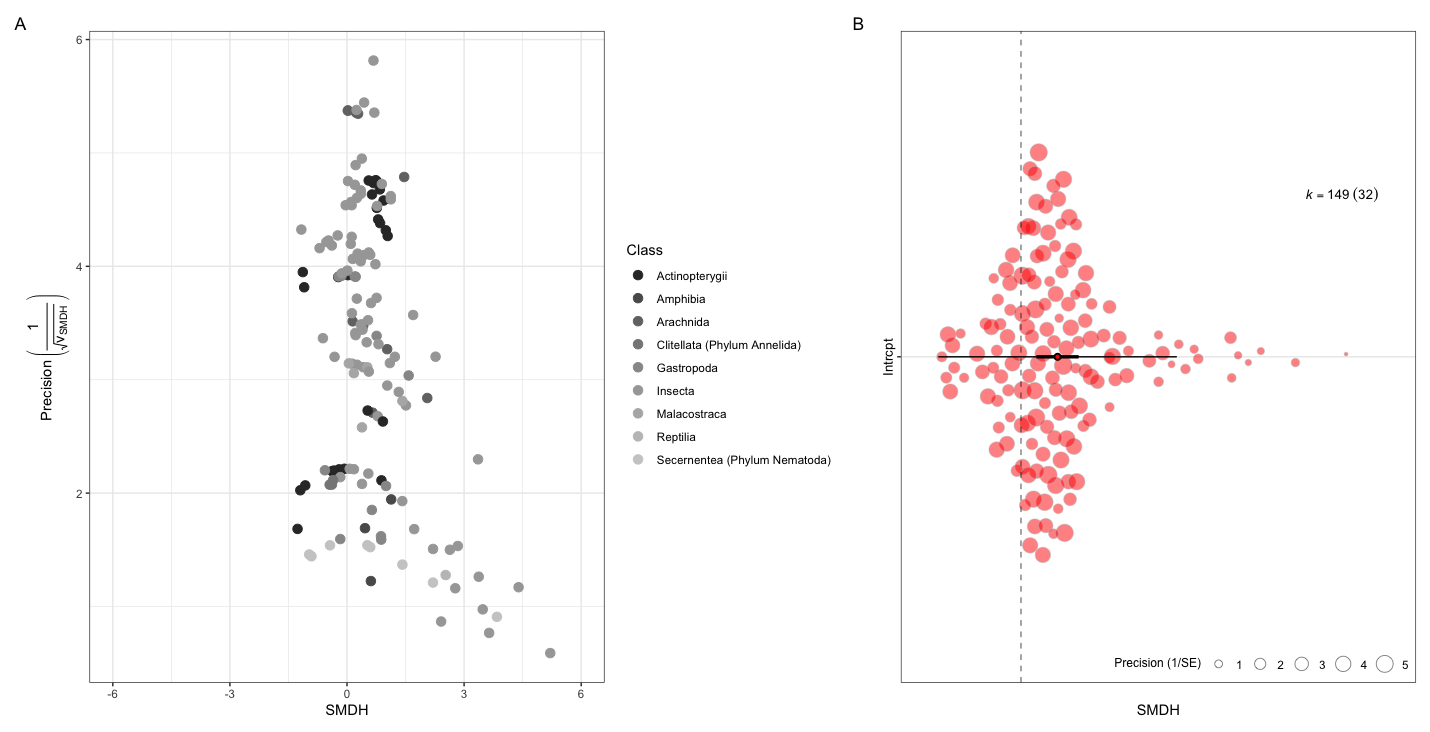

**Figure 1**. A) Funnel plot of effect size as a function of precision (i.e., inverse of sampling standard error). B) orchaRd plot of the overall meta-analytic mean effect size, 95% confidence intervals (thick black bars) and 95% prediction intervals (whiskers)
